# Viral lysis of a toxigenic diatom triggers a microbial response mimicking hastened senescence

**DOI:** 10.64898/2025.12.03.690933

**Authors:** Timotej Turk Dermastia, Tinkara Rupnik, Tinkara Tinta

## Abstract

Diatom blooms influence carbon cycling through organic matter production and its subsequent deposition or remineralization – processes that are all tightly mediated by interactions with the microbial community. Viruses, as integral part of microbial communities, are known to influence diatom bloom dynamics and can even terminate blooms. However, the three-way interactions between diatoms, their viruses and associated bacteria remain poorly resolved. In this study we examined how infection of the toxigenic diatom *Pseudo-nitzschia galaxiae* by its ssRNA virus PnGalRNAV reshapes host physiology, microbiome structure and organic-matter processing in non-axenic batch cultures. With an integrated transcriptomics and microscopy-based approach, we investigated the response of the bacterial community to dissolved organic matter (DOM) released by viral lysis of diatoms and observed a significant increase of Flavobacteriaceae (Bacteriodetes) suggesting a specialized role in utilizing DOM released due to viral lysis. Despite overwhelming viral RNA, bacterial metatranscriptomes revealed upregulation of polysaccharide-degradation associated genes, indicating active utilisation of virus-derived diatom glycans. Host transcripts showed broad repression of photosynthesis, silicon metabolism and core biosynthetic pathways, alongside induction of heat-shock and other stress-related genes, consistent with a senescence-like state. Our results demonstrate that PnGalRNAV infection speeds the termination of *P. galaxiae* growth, rapidly converting a productive diatom culture into a detrital DOM-rich environment that selects for specialised polysaccharide degraders and redirects carbon through the viral shunt. Infected cultures reached senescence much sooner than uninfected ones, suggesting that the ssRNA virus of *Pseudo-nitzschia galaxiae* can shorten bloom duration and accelerate nutrient recycling, with implications for coastal biogeochemistry. By integrating spatially resolved microscopy with community and metatranscriptomic profiling, this study links microbial composition, localization, and functional activity during diatom viral lysis. Our comprehensive approach can be extended to diverse diatom–virus systems in the future to better predict when viral outbreaks will favour recycling versus export of phytoplankton-derived carbon.

## INTRODUCTION

Phytoplankton, as a major eukaryotic, unicellular group, depend on the supporting bacterial community for growth and survival ^1^. The interactions between them occur in the phycosphere, the immediate surroundings of phytoplankton cells, where nutrient and chemical exchange between phytoplankton and bacteria takes place ^2^. Diatoms are important constituents of marine phytoplankton communities and are not exempt from interactions with bacteria ^3–5^. Various *Bacteroidetes, Gammaproteobacteria and Alphaproteobacterua* are often associated with diatom in situ and in cultures ^3,6–9^. Diatoms (and phytoplankton alike) are also affected by viruses. Viruses are an integral part of marine microbial communities ^10^. They outnumber bacterial cells by a factor of 10 and unicellular eukaryotes by a factor of 10^4^. Viruses infecting phytoplankton primary disturb the production of biomass by killing their hosts and transforming the particulate organic matter (POM) to the dissolved form (dissolved organic matter, DOM) fuelling the microbial loop, a process also known as the viral shunt ^11^. This process also leaves a distinct metabolic fingerprint in the derived DOM ^12,13^. Recent data also show that DOM derived from different viruses infecting the same host, elicits a very different response in the microbial growth ^14^. On the other hand, viruses can also enhance carbon export in certain conditions, which is known as the viral shuttle ^15^. This process mainly unfolds through the increase of aggregating potential of lysed or infected cells because of the production of sticky exudates ^16–18^. This process may also be influenced by bacteria and the ways they digest or transform organic matter produced by infected or dying cells ^19^. The interplay between phytoplankton, bacteria and viruses is however not well studied. Bacteria readily exploit the DOM released from lysed cells but may also have other indirect effects on phytoplankton. It has been shown for example that bacteria may confer resistance to certain viruses in *Chaetoceros tenuissiumus* cocultures ^5^. This finding adds to the understanding that phytoplankton have associated microbiomes that boost their fitness.

In this study we exposed a culture of *Pseudo-nitzschia galaxiae* to its lysing virus PnGalRNAV to better understand the response of the microbial community from the diatom culture to DOM released by lysed diatom cells as well as to the changed physiological state of infected but still living cells. In the experiment we monitored abundance of diatoms and specific bacterial populations via fluorescence *in situ* hybridization using epifluorescence microscopy andvirus abundance via qPCR, concentration of inorganic nutrients and DOM. We applied omic approaches (16S rRNA gene amplicon sequencing and metatranstcriptomics) to study dynamics microbial community structure and function

## METHODS

### Experimental setup

A non-axenic culture of *Pseudo-nitzschia galaxiae* (isolate Pn208, 28S accession number OR482970) was grown in 10-liter Duran® flask containing 8L of L1 medium ^20^. When the culture reached exponential phase of growth, determined by cell counts on a Fucks-Rosenthal counting chamber, the culture was distributed into six 1-liter pre 1N HCl-washed, MilliQ rinsed and autoclaved Erlenmayer flasks. The remaining 2L were kept for immediate analysis. Three of the 1-liter flasks were designated as controls and three were designated as treatments. Into the control vessels we added 30 ml of sterile medium. Into the treatment vessels we added 30 ml of a pre-harvested viral lysate of the PnGalRNAV obtained and stored as described in ^21^. The vessels were kept in a thermostatic chamber at 18°C on a shaker (400 rpm) and a 12:12 hour light:dark cycle with a light intensity of ∼100 µmol/photons/m^2^/s. The batch experiment was left to run for five days. We subsampled the batch cultures for diatom and bacterial abundance, abundance of specific bacterial populations using specific FISH probes, nutrient and DOC concentrations, and virus concentrations for qPCR. Briefly, 60ml from the experimental cultures was subsampled using acid-cleaned and combusted glass pipettes. 10ml were aliquoted for bacteria, diatom and virus abundance and composition counts. 15ml of the subsample was filtered through combusted 25 mm GF/F filters (Whatman) and stored in a 15 ml plastic tubes for nutrient analysis, while the rest was kept in acid cleaned and combusted glass vials for DOC analysis. The filters were stored at -20°C. Samples for 16S community profiling and metatranscriptomics were collected in the initial batch culture and five days post infection (DPI). At five DPI there was ∼700ml of culture left in the flasks prior to filtration to collect nucleic acids, which is within the recommended ⅔ of the initial volume to ensure representability. After filtration the remaining volume was ∼500ml. We decided to keep the experiment going although representability was not ensured and subsampled again nine DPI for all parameters except for nutrients and DOC.

### Diatom and bacterial counts

For living diatoms, one millilitre of the culture was collected and fixed in 2.5% formaldehyde (final concentration) and stored at room temperature on each day of the experiment (including Day 9). The samples were counted within two weeks after the experiment on Fucks-Rosenthal and Thoma counting chambers under the Axio Imager Z2 (Zeiss) epi-fluorescence microscope (EFM). For bacteria and dead diatoms, 1mL of sample was collected from every experimental bottle at every time point and fixed with 2% formaldehyde (final concentration) and stored at -80 °C until further processed, mounted with DAPI (Ex/Em = 358/461 nm) and examined with EFM at 1250x magnification, as described previously ^22^ We partitioned the bacterial counts into free-living bacteria (FLB) and diatom attached bacteria (DAB). Living diatom cells were identified by the presence of an intact DAPI-stained nucleus and two chloroplasts (Supplementary Figure 1A,C,E,G,I). In addition, we quantified diatom detritus–associated bacteria, corresponding to bacteria attached to dead or lysed diatom cells, which were recognized by a visible frustule or by chloroplasts in the absence of a DAPI-stained nucleus (Supplementary Figure 1B,D,F,H,J). This definition also corresponded to our identification and enumeration of dead diatom cells.

### Fluorescence in situ hybridization

The abundance of specific bacterial populations were determined by fluorescence in situ hybridization (FISH) using specific oligonucleotide probes labelled with fluorophore Cy3 or FITC at the 5’-end (Biomers) (Supplementary Table S1). The specific bacterial populations were selected based on the relative abundances of bacteria from metatrascriptomes/16S rRNA amplicon sequencing data. We applied a modified version of the FISH method as before ^23^.

### Virus quantification

To quantify the virus, we developed a TaqMan qPCR assay following MIQE guidelines ^24^. The assay is described in Turk Dermastia et al. (2024).

### Dissolved organic matter and nutrient measurements

Samples for dissolved organic carbon (DOC),total dissolved nitrogen (TDN) and inorganic nutrients were analysed as we describe in detail in our previous experiments ^22^.

### DNA and RNA analysis of diatoms and bacteria

The remainder of the culture from the beginning of the experiment was immediately filtered to assess the initial state of the culture: 500 ml onto 1.2 µm cellulose filters in duplicate for metatranscriptomis of diatoms and associated microbial communities; 200 ml onto 0.22 µm polycarbonate filters for 16S bacterial community analysis and onto 1.2 µm cellulose filters for metatranscriptomics at start of the experiments, on Day 5 and at the end of the experiment (Day 9) from control and virus treatments (Supplementary Table 2). All filters were instantly frozen and stored at -80°C until further analysed. DNA from bacterial samples was extracted using the DNeasy PowerWater Kit (Qiagen, Hilden) following the manufacturers guidelines. 16S was amplified using primers from ^25^. Library preparation for Illumina HiSeq 2x250bp and sequencing with a depth of 100.000 tags per sample were performed by Novogene Ltd (Cambridge, UK).

Filters for RNA extraction were submerged in TriZol reagent (Zymno Research), bashing beads were added, and the mixture was vortexed vigorously for a minute. The mixture was then flash frozen, thawed and vortexed again for a minute. This was repeated 3 times, before proceeding with extraction according to the manufacturer’s protocol using the Direct-zol RNA Miniprep kit (Zymno Research). The RNA concentration and integrity was measured with a HS RNA assay on the Tapestation (Agilent). Sequencing was performed on a NovaSeq X Plus Series (PE150) by Novogene Ltd.

### Bioinformatic analysis

#### 16S amplicon analysis

The outputs from 16S sequencing were, quality-checked, merged and trimmed using the *DADA2* standard pipeline and compared to the SILVA 99% 16S database (v138) for community composition inference. Alpha and beta diversity analyses were conducted in *phyloseq* ^26^ and *vegan* packages ^27^.

#### Microbial metatranscriptomes

The entire pipeline can be found in Supplementary Methods. Scripts are available through https://github.com/Timzladen/PnGal_transcriptome.

## RESULTS

### Host population dynamics

#### Viral attack and organic matter release

Both the control and virus treatments showed an initial increase in diatom cell numbers towards Day 3 of the experiment, followed by a quick collapse of the culture in virus-infected treatments, and a more gradual decrease in cell numbers in the non-infected controls (Figure 1A). The collapse coincided with the stationary phase of diatom population growth in both treatments, however in the control treatment the collapse of population was delayed for a few days. Concurrently dead diatoms increased in both treatments, but the change was more pronounced in virus treatments (Figure 1B). The viral copy number followed the pattern of diatom population dynamics and interestingly also decreased about 6-fold to initial inoculum values as the diatom culture collapsed (Figure 1C). However, almost 10^4^ copies of the viral RdRP gene remained detected also after this collapse. Virus-treated cultures visibly discoloured by Day 5, consistent with rapid lysis, while controls retained the brown pigmentation of healthy diatom cultures. (Supplementary Figure 2).

**Figure 1.**
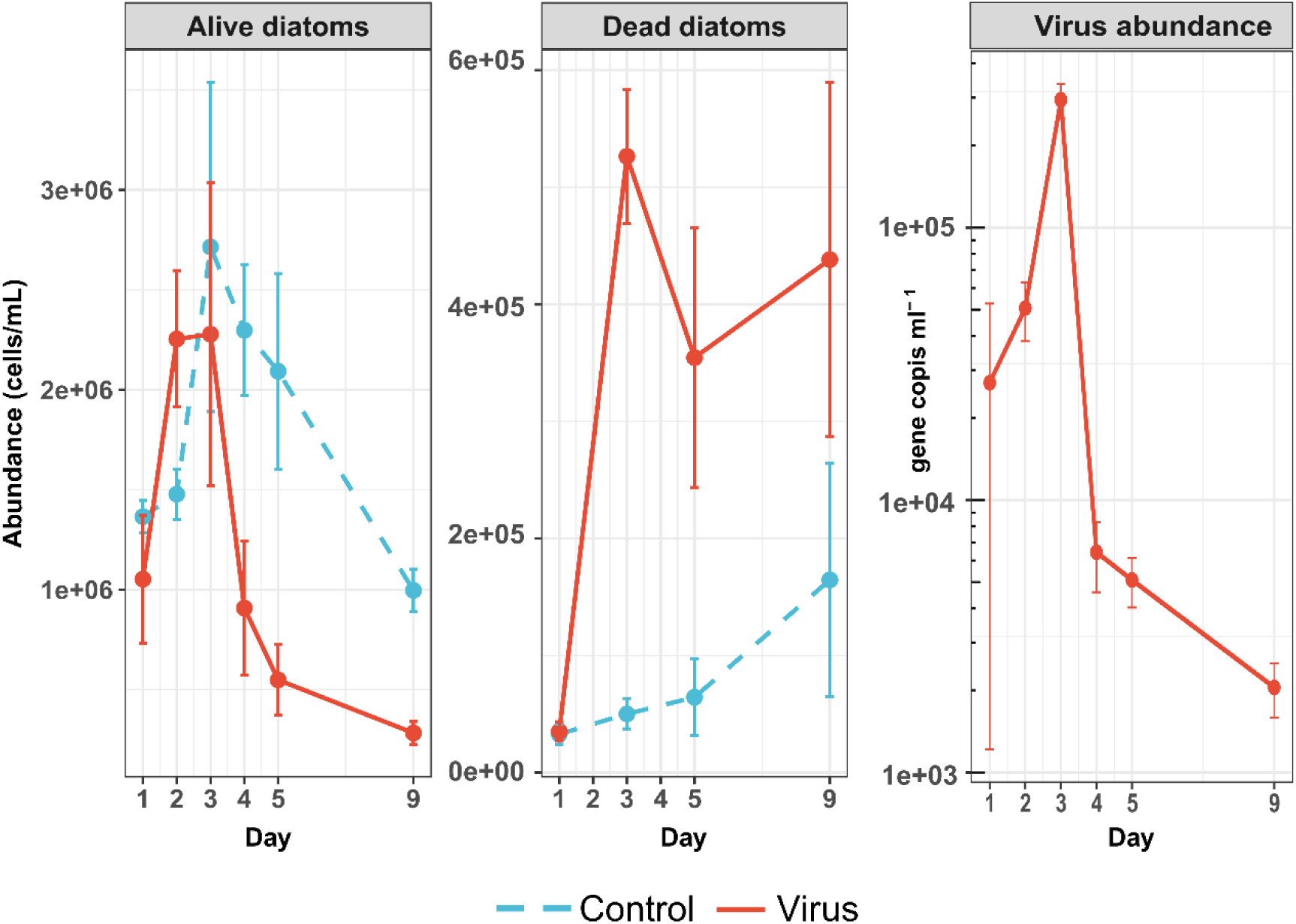
Host-virus dynamics obtained through different methods. Live diatoms were counted in Fuchs-Rosenthal chambers in living conditions. Dead diatoms were inferred from DAPI staining (cells with stained chloroplasts but not nuclei). Virus concentrations were obtained by qPCR.

The DOC concentration increased almost 2-fold from the start of the experiment onwards in virus-infected cultures and 1.5-fold in controls, as both the control and virus infected cultures started to decline (Supplementary Figure 3). DOC accumulation in virus-treated samples became significantly greater than in controls from Day 4 onwards (*t=*4.804, *p < 0.001*), potentially reflecting increased organic matter release due to diatom lysis. DOC was highest on Day 5 in two virus infected flasks, reaching 586 µM and 583 µM in Virus1 and Virus2, respectively. The diatoms in both the virus treatment and controls depleted nitrate, silicate and phosphate without significant differences between the treatments (Supplementary Figure 3). On the other hand, there was a stronger buildup of ammonium in the virus treatments. The difference was not overall statistically significant but after estimating for marginal means we saw a significant change in Day 5 (*t=*3.479, *p < 0.01*).

### Dynamics of microbial community

#### Structure of microbial community based on amplicon sequencing of bacterial 16S rRNA genes

Amplicon sequencing detected 333 ASVs, with low diversity across treatments (Shannon index ≤ 2.5; species richness ∼25, Supplementary Figure 4). Principal coordinate analysis (PCoA) showed strong separation of initial samples from both treatments, and convergence of control with virus-treated samples, indicating that senescent controls shifted toward a virus-like community state (Figure 3A). ANOSIM supported significant overall group differences despite non-significant pairwise tests. Both treatments exhibited strong temporal restructuring. *Marinobacter* (Gammaproteobacteria) declined sharply (from 60% to 5%) from early dominance to low abundance by Day 5, while Bacteroidetes—primarily *Polaribacter*—expanded dramatically, from 20% reaching ∼70% relative abundance in virus-treated flasks and ∼50% in control. Other Bacteroidetes genera mostly from the Flavobactericeae family were *Fluviicola* (Figure 3B, Supplementary Figures 5) in the virus-treated flasks and *Flegellimonas* (Flavobacteriaceae, Bacteroidetes) in the controls. The latter also showed an increased relative abundance in control flasks as the experiment progressed (Supplementary Figure 5C). Control flasks also showed increases in unclassified Rhodobacteriaceae (Alphaproteobacteria) (Supplementary Figures 5A and 5B). DESeq2 confirmed significant enrichment of *Polaribacter* in virus treatments and Rhodobacteriaceae in controls (Figure 3E, Supplementary Table 3).

**Figure 2.**
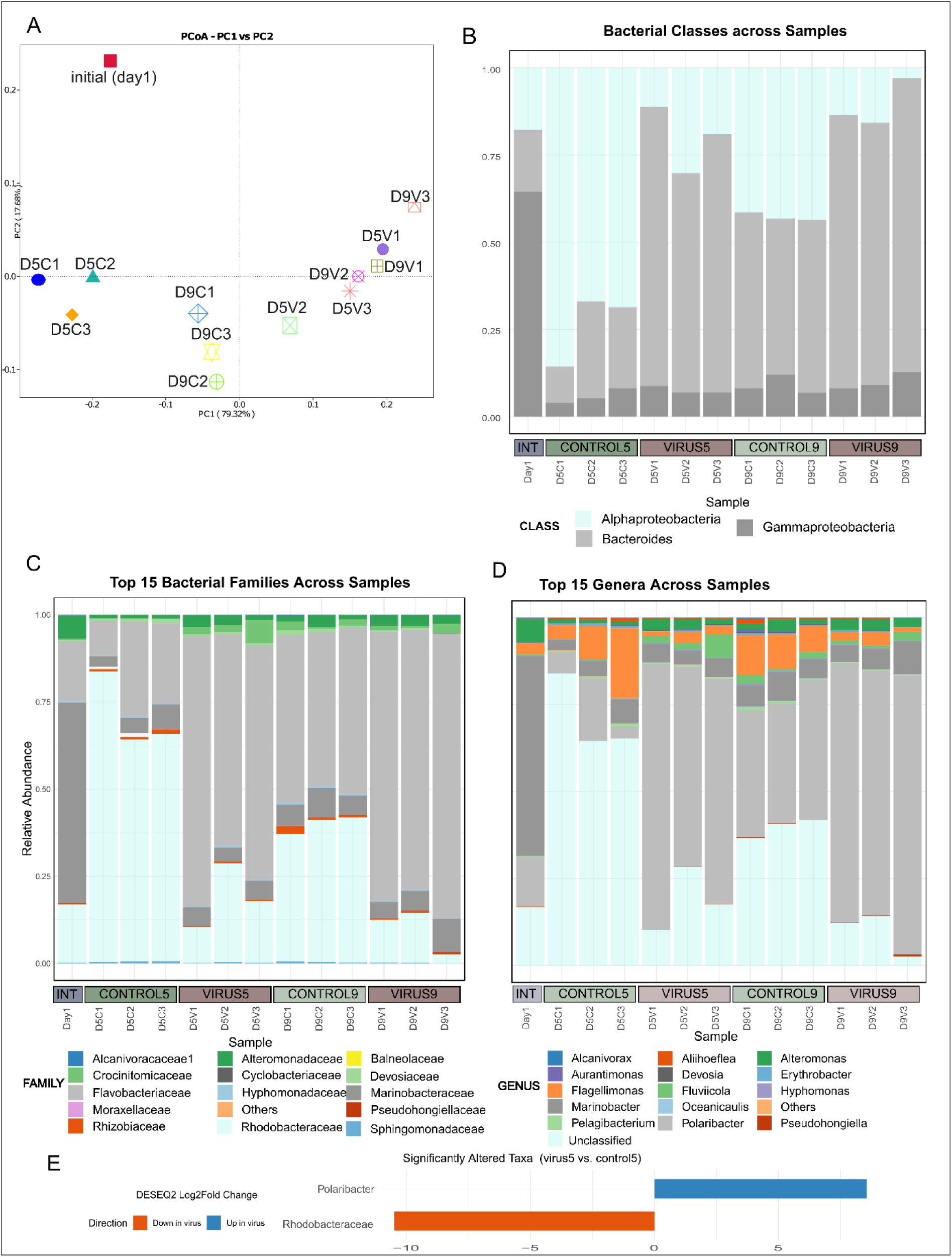
Bacterial genus taxonomy inferred from 16S amplicon sequencing. See supplementary Figure for complete representation. (A). PCoA with weighted unifrac distances. (B) Relative abundances of bacterial classes. (C) Relative abundances of Top 15 bacterial families. (D) Relative abundances of top 15 bacterial genera. (E) Significantly altered taxa from DESEq2 analysis and their log2fold changes between controls at Day 5 (control 5) and infected cultures at Day 5 (virus5).

**Figure 3.**
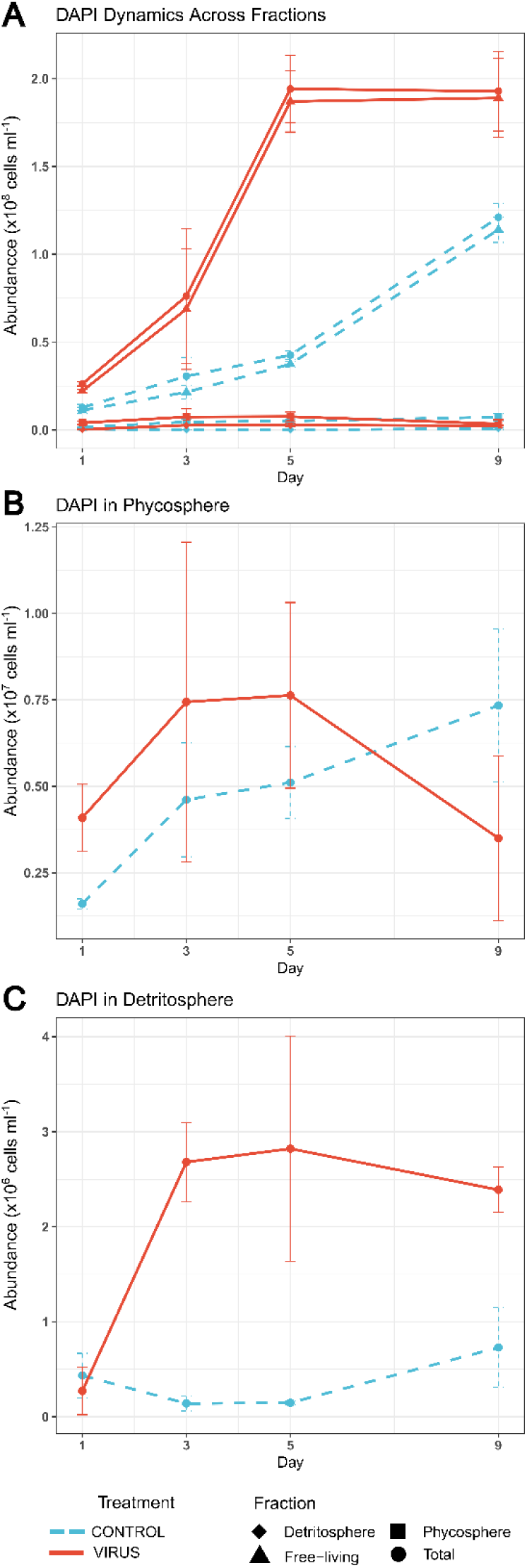
Dynamics of the microbial population from DAPi staining in different fractions (A). Close up for scaling of phycosphere associated microbes (B), and detritosphere associated microbes (C). Error bars are standard errors.

#### Population dynamics of specific microbial populations via microscopy-based approach

Total microbial abundance (DAPI-stained cells) rose tenfold by Day 5 and remained stable through Day 9 in virus treatments (Figure 3A). Controls increased more gradually, reaching ∼1.5-fold lower abundances by Day 9 as compared to virus treatments. EUB-positive cells constituted ∼88% of DAPI-stained microbes in controls and ∼93% in virus-treated cultures (Supplementary Table 4). The abundance of phycosphere-associated microbes was initially similar in both treatments, then (Figure 3A) increased steadily in controls (∼4.6-fold by Day 9), whereas in virus treatments rose modestly to Day 5 (∼1.8-fold), then decreased as diatoms collapsed (Figure 3B). The detritosphere fraction increased profoundly in the virus treatments and remained constant in controls (Figure 3C). Attached bacteria per live diatom increased from ∼2.5 (control) and ∼3.3 (virus) on Day 1 to ∼10 in controls and ∼30 in virus treatments by Day 5, followed by a decline in virus cultures (Supplementary Figure 6A). Attached bacteria per dead diatom declined from ∼13 to ∼3 per cell by Day 3 and remained constant throughout the experiment in controls (Supplementary Figure 6B), while in the detritosphere microbial load remained stable at ∼5–8 bacteria per cell in the virus treatments.

Bacteroidetes (CF319a FISH probe) drove most of the microbial community increase in both treatments (Figure 4 and 5). In virus treatments, they rose ∼6.5-fold by Day 3 and dominated the microbial community, thereafter, reaching ∼ 70% and only ∼ 49% of DAPI-stained cells in the control by Day 9. Alphaproteobacteria increased more in controls than in virus treatments (∼1.9-fold difference), while Gammaproteobacteria declined in both, from ∼ 45% to to ∼13% in control) and from 63% to ∼9% in virus treatment from start until the end of the experiment. b Free-living fractions showed comparable trends.

**Figure 4.**
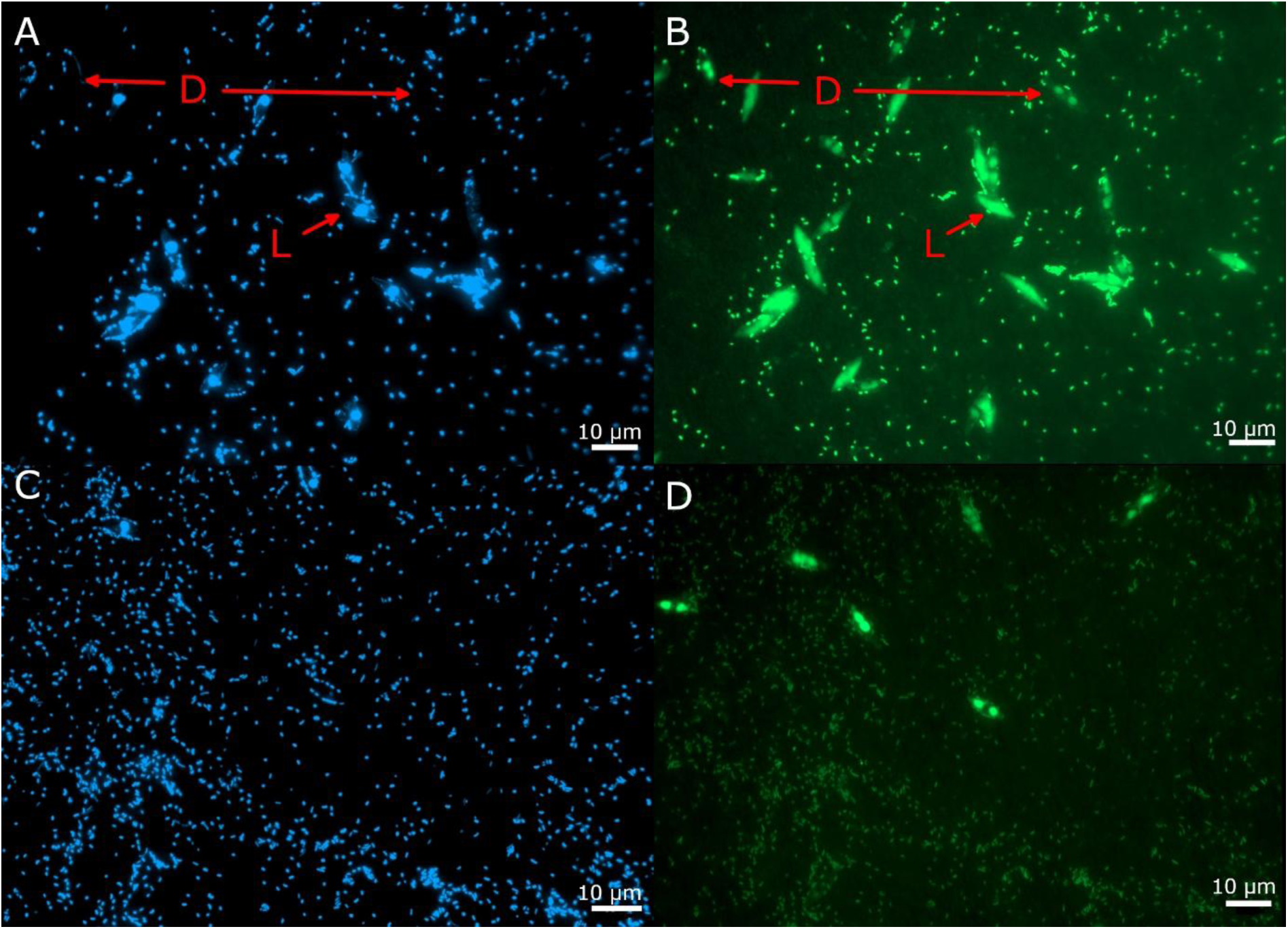
DAPI (A) and CFB (B) stained sample of infected diatoms 3 days post infection (A and B) and 5 days post infection (C and D). Dead diatoms (D) are identified by a missing DAPI stain but visible signal with the other probe. In this case chloroplasts are clearly visible. Bacteria attached to such cells were counted as detritosphere associated. In contrast, living cells (L) have clearly stained nuclei with DAPI. Bacterial cells attached to such cells were counted as phycosphere associated.

**Figure 5.**
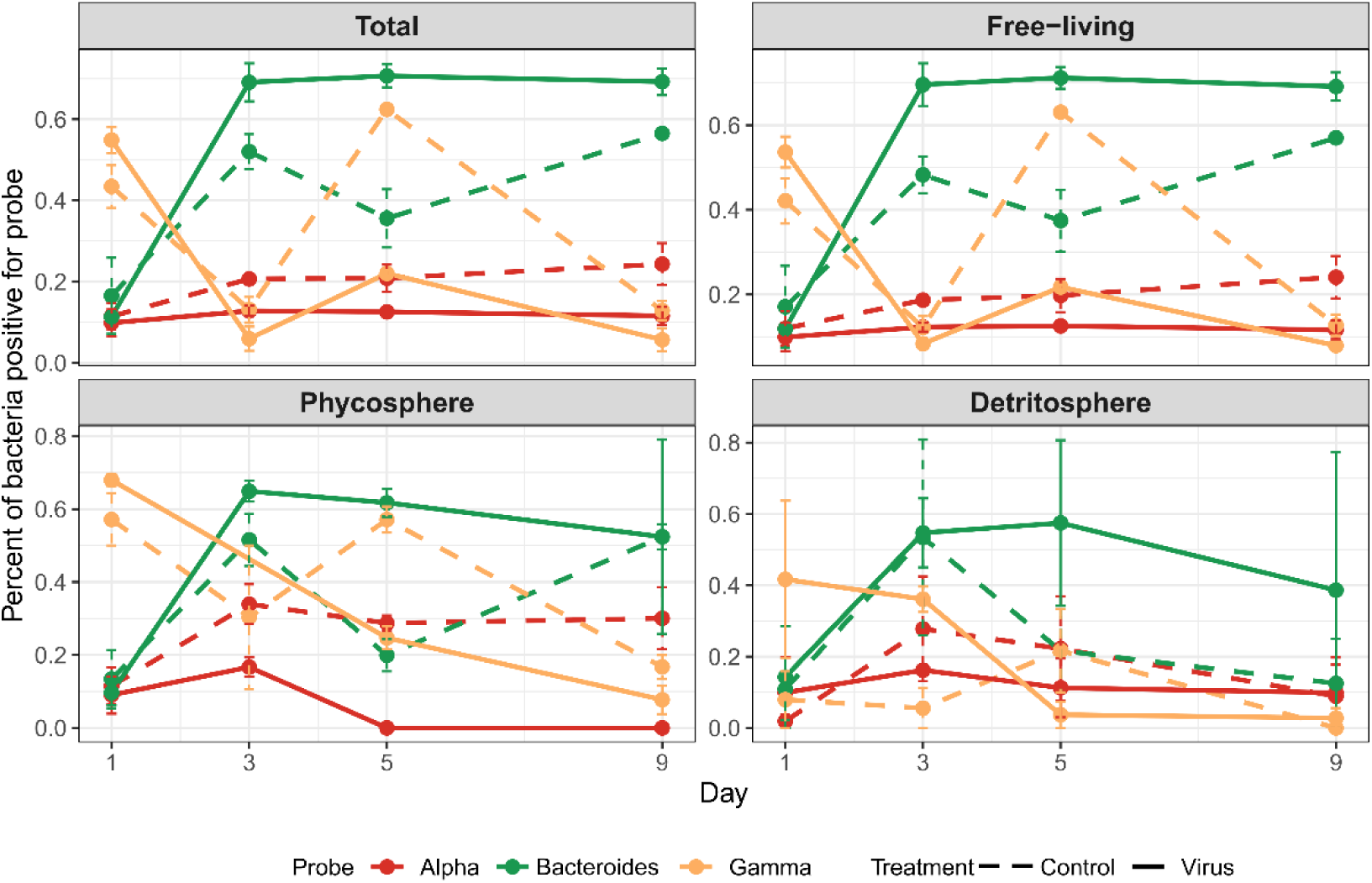
Dynamics of specific bacterial populations within total microbial community, free-living population, phycosphere- and detritosphere-associated community in control vs virus treatments over time. Alpha-Alphaproteobacteria; Gamma-Gammaproteobacteria.

Bacteroidetes also dominated the phycosphere (i.e., microbes attached to live diatoms, Figure 5) in virus treatments after Day 3, reaching ∼80% by Day 9, while in the controls they represented ∼20–50% of DAPI-stained cells. Gamma- and Alphaproteobacteria were consistently depleted in the phycosphere relative to the bulk community. Detritosphere patterns were more pronounced: Bacteroidetes reached ∼77% of cells associated with dead diatoms in virus treatments by Day 9, compared to ∼37% in controls. Gammaproteobacteria associated with dead diatoms dropped from ∼70% to ∼7% in virus treatments but remained relatively stable in controls. Alphaproteobacteria showed a transient peak on Day 3 in virus treatments before declining, with similar pattern, but slightly higher abundance in the detritosphere of controls.

By Day 5, attached Bacteroidetes reached 18.6 cells per diatom in virus-treated cultures versus 2.1 in controls (ANOVA: F = 5.589, p = 0.024), though replicate variability limited day-specific resolution.

### Insights into metabolism of microbial populations

#### Metatranscriptome sequencing results

Sequencing generated ∼80-100M reads per sample. The number of reads belonging to the host, bacterial community and viruses varied depending on the condition from which the sample was obtained. Viral-infected samples five DPI were already heavily dominated by viral RNA, representing 75% of the metatranscriptome, potentially limiting the metatranscriptome insights into the host and microbial functions (Supplementary Table 5). In Day 5 controls, 50–60% of TPM belonged to bacteria, compared to ∼2% in virus-treated samples.

#### Transcript-level bacterial taxonomy

The taxonomical composition on the transcript level displayed similarity to the taxonomy obtained with 16S sequencing with some notable differences (Supplementary Figure 7). For example, *Flagellimonas (Bacteroidetes)* which took up large relative abundances in 16S was scarce in the transcript taxonomy. On the other hand, *Marinobacter (Alphaproteobacteria), Polaribacter (Bacteroidetes),* and *Alteromonas (Gammaproteobacteria)* showed congruent patterns, with *Polaribacter* increasing in relative abundance through the course of the experiment, while *Marinobacter* decreased and *Alteromonas* remained constant without major changes. Some other bacterial genera emerged in this analysis, however sticking to the 16S taxonomy is likely a more accurate representation of the community.

#### Broad expression patterns

Despite lower bacterial transcript counts in virus samples, functional profiles were conserved across initial, control, and virus conditions (Figure 6, Supplementary Figure 8). Enriched COG categories included energy production/conversion (C), translation (J), cell wall biogenesis (M), and amino acid metabolism (E). A substantial portion of genes fell into unknown function (S). Taxonomically, functional contributions shifted from *Marinobacter* dominance in initial samples to Flavobacteriaceae dominance later. In virus-treated cultures, *Polaribacter* replaced *Marinobacter* across most COG categories, notably dominating defence mechanisms (COG V) and carbohydrate-related pathways. Some COGs showed distinctive contributors: *Maricauda* dominated energy production in virus samples, while *Crocinitomix* and *Fluviicola* contributed to specific ion transport and trafficking functions.

**Figure 6.**
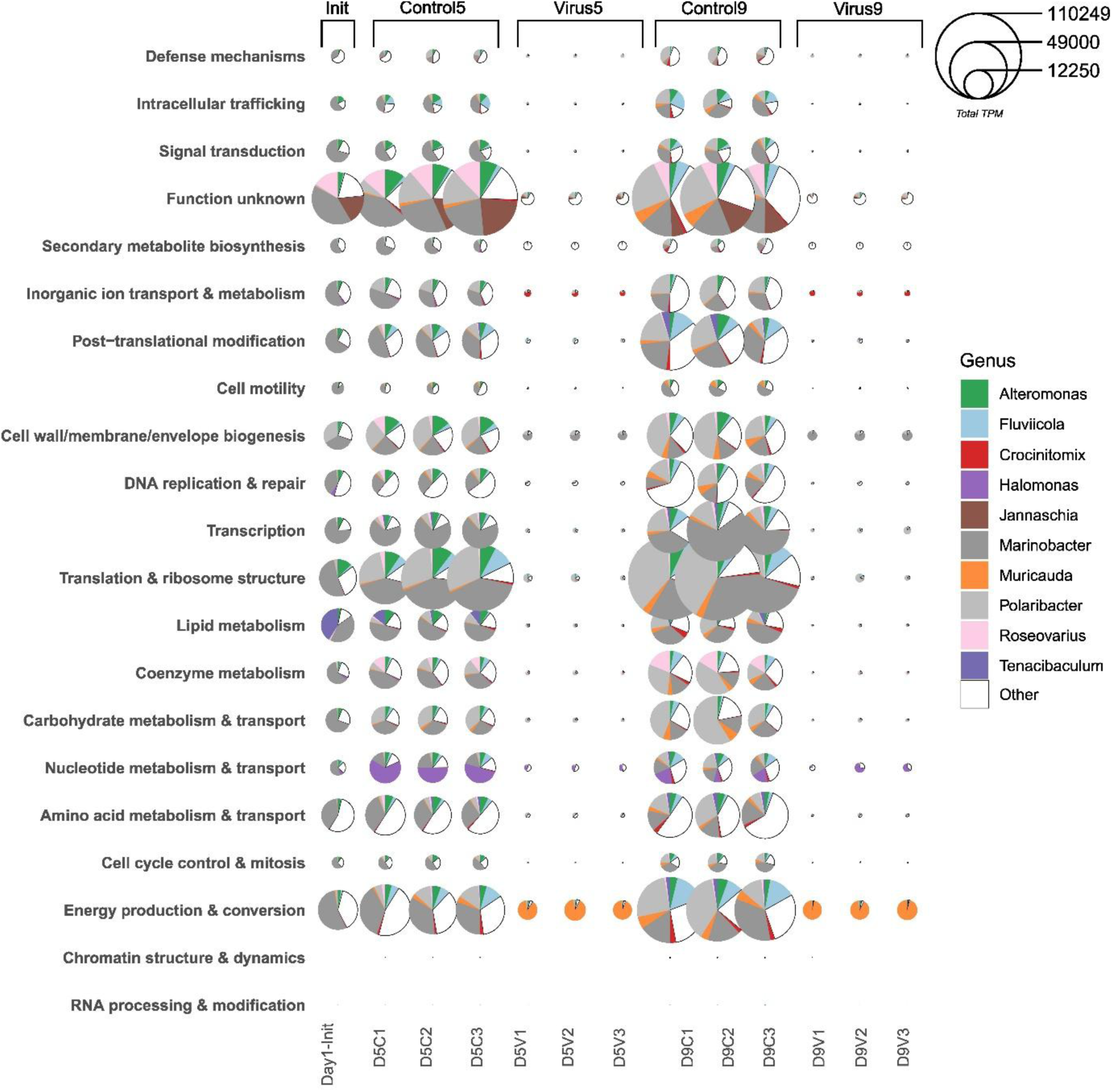
Absolute TPM of COGs from different genera. For relative representation see Supplementary Figure 8.

#### Differentially expressed bacterial genes

A total of 128 bacterial genes were differentially expressed between infected and control cultures at Day 5 (Supplementary Figure 9). Most were downregulated in virus treatments due to community restructuring and transcriptional suppression. Virus-unique upregulated categories included post-translational modification (O), nucleotide metabolism (F), and carbohydrate transport/metabolism (G), highlighting increased processing of diatom-derived DOM. Consistent with community turnover, upregulated genes were predominantly assigned to *Polaribacter* and other Flavobacteriia (Figure 7, Supplementary Table 6). These bacteria were actively using diatom-derived carbohydrates as a source of carbon, as indicated from upregulation of genes such as *cellobiose phosophorylase, TonB, transglycosylase*. Several metabolic and regulatory signatures were also enriched: carbohydrate metabolism enzymes, enzymes associated with oxidative stress response, lipid and aromatic compound metabolism, nitrogen metabolism (glutamate synthase, GAD proteins), and motility/stress-related proteins (heat shock proteins). Genes linked to transcriptional regulation (RNA polymerase subunits, restriction-modification systems, Sec translocase ATPases) were also enriched, indicating broad reprogramming of bacterial metabolism. Only one DE gene was found when comparing infected samples at Day 5 and controls at Day 9 (Supplementary Table 7), reflecting converging community states but also compositional bias. Analysis without transcript clustering returned less DE genes where some were unique, primarily additional *Polaribacter* enzymes like *ATP-dependent exoDNases*, and *nicotinic-acid mononucleotidases*, and a bigger suite of carbohydrate degrading enzymes (Supplementary Table 8)— suggesting broader participation in carbohydrate and nucleic acid degradation and salvage pathways.

**Figure 7.**
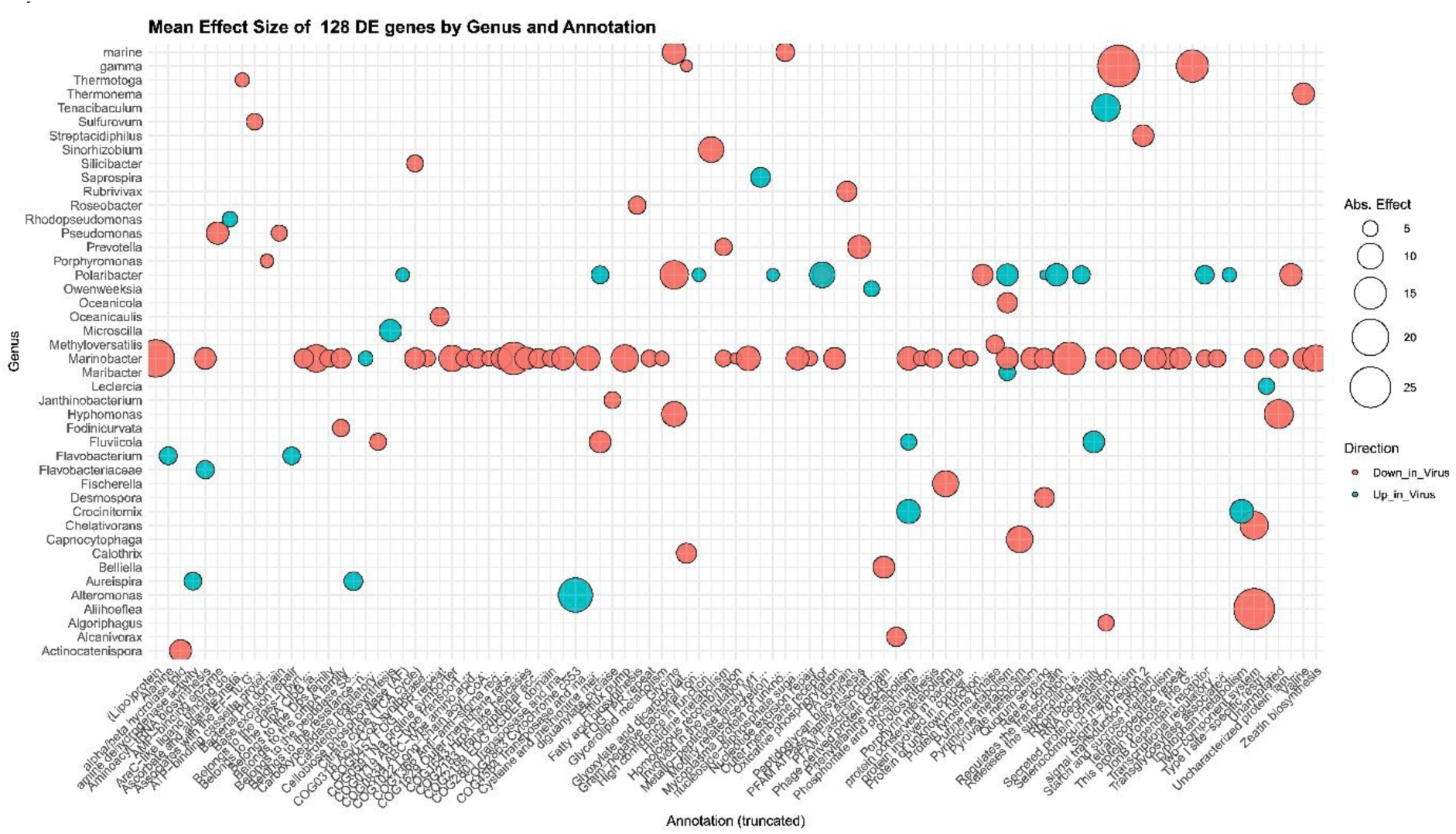
DE expressed genes and their taxonomic origin (ALDEx2) between infected samples and controls at Day 5.

#### Bacterial CAZymes

Marker genes for six key glycan types — alginate, β-glucan (laminarin), β-glucan, β-mannan, chitin, exo-polysaccharides, host glycans and xylan— were selected based on literature-defined combinations of GHs, PLs, CBMs, and relevant transporters (SusD and TonB). For each substrate, functional completeness was calculated as the proportion of CAZy gene families detected per sample (TPM > 0.1) relative to the total number of distinct families observed for that substrate across all samples (Supplementary Table 9). Of ∼1600 detected CAZy genes, glycosyltransferases and glycoside hydrolases showed the clearest virus-associated shifts. Substrate profiling revealed lower total CAZyme TPM in virus-treated samples—driven by compositional dilution—but relative expression of many families increased in Flavobacteria, particularly *Polaribacter* (Figure 8). β-glucan (laminarin)-related genes showed the highest completeness after host glycans, increasing in control samples and remaining functionally represented in virus treatments despite lower total TPM. Host glycan-associated CAZymes (15 families) were abundant in initial and control samples but reduced in virus cultures; however, several Flavobacteria maintained relative expression under infection. Other substrates were represented by fewer families: alginate (3), β-mannan (2), chitin (5), exopolysaccharides (2), and xylan (2). Their completeness profiles were relatively uniform across samples, though expression was often low. A notable exception was *Flaviramulus*, whose alginate lyase genes showed elevated absolute and relative expression specifically in virus-infected samples, accompanied by borderline-significant differential expression, accompanied by borderline-significant differential expression (effect > 7, p raw < 0.01, FDR > 0.1). Overall, β-glucan, β-mannan, chitin, exopolysaccharide, and host glycan genes were broadly distributed taxonomically between different Flavobacteria, while alginate and xylan degradation appeared to be genus-specific. TonB/SusD transporters—key markers of glycan scavenging—were significantly upregulated infected cultures at Days 5 and 9 relative to control cultures at Day 5 (Supplementary Figure 10), but not relative to controls at Day 9 — consistent with the high TPM values present in control9.

**Figure 8.**
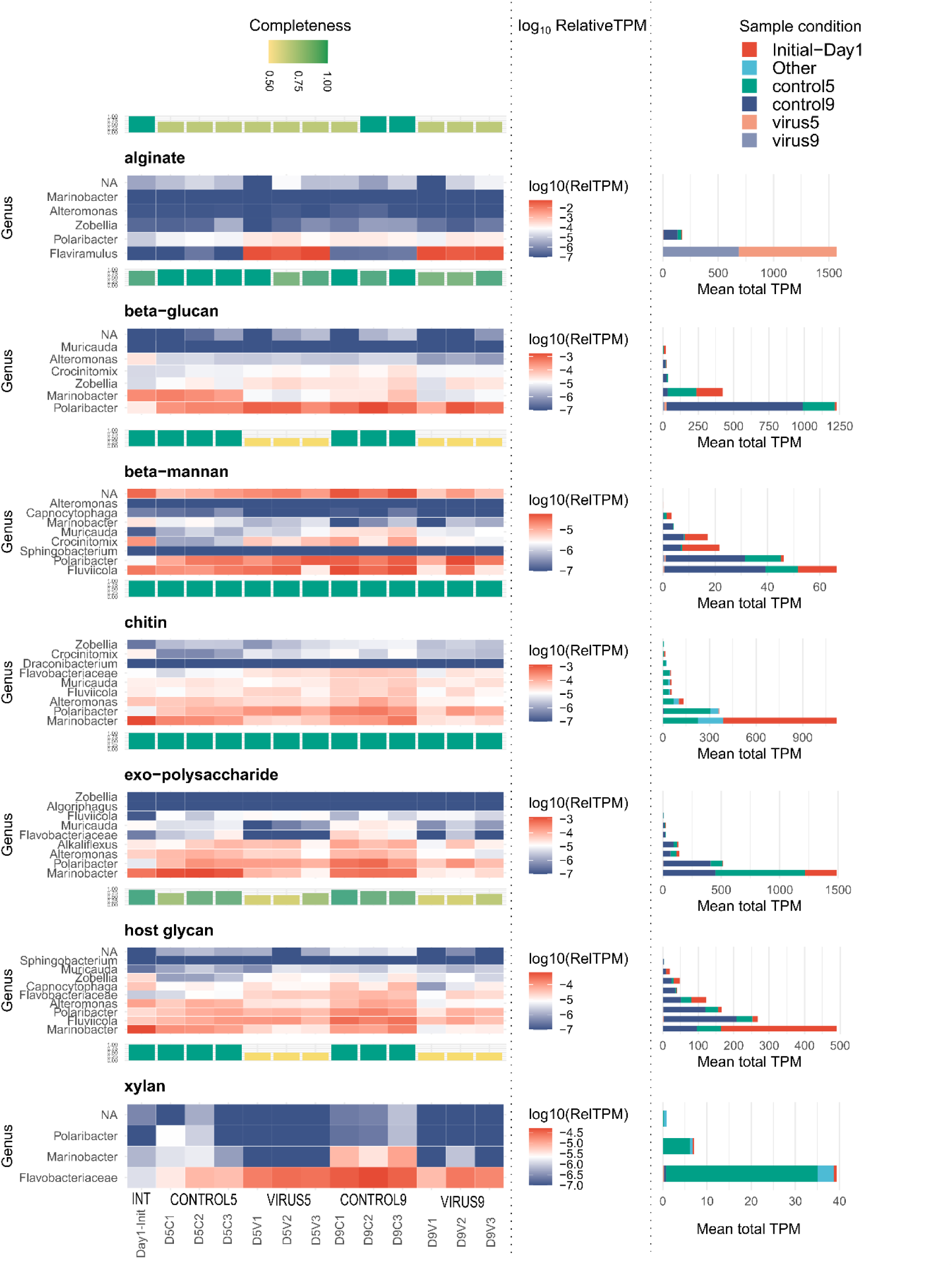
Heatmaps represent glucan pathway expression (log10(relativeTPM), calculated as transcriptTPM/total sample TPM). Barcharts above show the substrate degradation CAZy family completeness, calculated as the proportion of CAZy families detected (TPM>0.1) versus the total number of CAZy families present in all samples. Plots on the right show the average total TPMs of all genes for each condition. Control/Virus5: Day 5; Control/Virus9: Day9, Int: initial condition (Day1).

#### Host-transcriptomes

The number of host-derived transcripts was low in virus-treated samples (∼2000 TPM), as viral RNA dominated the metatranscriptome. This imbalance constrained our ability to resolve host expression patterns, even though sequencing depth was high. BUSCO analysis reflected this skew: initial samples showed full recovery of conserved orthologs, controls showed partial recovery, and virus-treated samples showed markedly reduced completeness (Supplementary Figure 11). Consistent with this, ALDEx2—which accounts for compositional bias—identified only a single significantly differentially expressed host transcript between infected and controls at Day 5. To obtain qualitative insight despite these limitations, we also evaluated the data with DESeq2. Although the number of confidently recovered genes was small, virus-treated samples showed a general reduction in transcripts associated with key metabolic and biosynthetic processes, including inorganic ion transport, amino acid and coenzyme metabolism, intracellular trafficking, and cytoskeletal organization (Supplementary Figure 12). In contrast, several genes linked to protein turnover and chaperone activity (COG category O), particularly members of the HSP70 and HSP90 families, were more abundant in virus-treated samples. Heatmaps of TPM-normalized expression confirmed that these stress-related transcripts increased in multiple infected cultures, while samples collected at 9 DPI showed an overall decline in host transcript abundance consistent with advanced infection and culture collapse (Supplementary Figure 13). A similar stress-related signature was also observed in the senescent control9 sample. Furthermore, silicon transporters were reduced in infected cultures at Day 5 (Supplementary Figure 14), consistent with the general decline in biosynthetic activity.

WGCNA modules supported these trends (Supplementary Figure 15, Supplementary Tables 10 and 11). A large photosynthesis/translational module (turquoise) was strongly downregulated under infection. A stress-associated module (black) was positively correlated with viral treatments although the majority of transcripts had bacterial annotations. Two additional modules (*brown* and *magenta*) displayed transitional behaviour: positive correlation in early infected samples but negative correlation in late-stage infection and senescence, suggesting transient host adjustments that dissipated as viral takeover progressed.

Together, the host transcriptome results point to a consistent but coarse pattern: strong viral RNA dominance, suppression of core metabolic processes, and activation of a limited stress-response program, with finer-grained host regulatory dynamics obscured by the overwhelming abundance of viral transcripts.

## DISCUSSION

### Viral takeover and *accelerated* senescence of the diatom host

This study is one of the first to examine the transcriptional and compositional response of the diatom associated microbiome to viral infection of its host. In the *Pseudo-nitzschia galaxiae*–PnGalRNAV type-II system, viral infection led to rapid culture collapse within several days, consistent with previous characterization of this virus ^21^. Although viral load in this experiment was estimated by qPCR rather than by MPN assays, the inferred inoculum (∼10^5^–10^6^ viruses mL⁻^1^) was broadly comparable to earlier measurements based on MPN (10^7^–10^8^ mL⁻^1^ in stock lysates, unpublished data). The subsequent decline in qPCR signal after Day 3, despite the dominance of viral transcripts at Days 5 and 9, likely reflects technical artefacts of the filtration step—where viruses entrapped in cellular debris or extracellular matrix were removed— or the potential sequence divergence within the highly variable capsid protein gene targeted by the assay, a common characteristic in ssRNA viruses ^28^. Because sampling for metatranscriptomics was done on Day 5, when most cells were already lysing, our metatranscriptomic analysis captured the late stages of infection rather than the onset of host responses. By Day 9, both infected and control cultures had reached advanced physiological decline, displaying similar bacterial community structures, repressed silicon metabolism, and elevated stress-related transcripts. This convergence suggests that viral infection may accelerate processes otherwise occurring during natural senescence. Nutrient limitation and silicate depletion are known to trigger oxidative stress and programmed cell death in diatoms, producing apoptosis-like morphology ^29^. Nutrient depletion and increase of ammonium was significantly higher in infected cultures than in controls. The observed increase in the abundance of bacteria in the later samples could indicate that accumulated ammonium results from bacterial degradation of proteinaceous fraction of organic matter released during diatom cell lysis via the ammonification process ^30^. The infection thus appeared to telescope normal senescence into a rapid lytic event followed by bacterial growth—consistent with the “viral shunt,” whereby lysis redirects particulate organic matter into dissolved organic pools available for heterotrophic bacteria uptake ^31–33^. In our system, the virus-infected culture at five DPI phenocopied a nine-day-old control culture, highlighting the capacity of ssRNA viruses to drive premature bloom collapse and nutrient recycling.

### Bacterial exploitation of viral lysis products

Viral infection of *Pseudo-nitzschia* cells induced cell lysis, which resulted in release of organic substrates from decaying diatom cells, evident from increase of DOC concentrations. The DOM was in turn taken up by specific bacteria in the resident microbial community. These findings demonstrate that PnGalRNAV-mediated lysis produced particulate and dissolved organic matter readily utilized by heterotrophic bacteria

The phycosphere is a chemically structured boundary layer in which algal exudates and EPS regulate bacterial colonization, attachment strength, and the balance between mutualism and antagonism^34^. The notable decrease in phycosphere-associated microbes in virus-treated cultures after Day 5 suggests that viral infection destabilizes the phycosphere by interfering with diatom–bacteria interactions. Our findings indicate that infected or recently lysed diatom cells initially function as hotspots for bacterial attachment – as reflected by the increase in bacteria attached to dead diatoms between Days 3 and 5, followed by a decline of detritosphere-associated cells and a concurrent rise in the free-living fraction. Thus, these cells are not stably colonized, likely because viral lysis accelerates structural breakdown, producing smaller, less cohesive particles from which attached bacteria readily detach and transition to the free-living phase. This reflects models of marine snow and phytoplankton aggregate microbiology, which emphasize that aggregates act as transient nutrient hotspots subject to intense colonization, rapid turnover, and high rates of detachment driven by particle disaggregation and fluid shear^35^. Moreover, virus-infected and non-infected dead diatoms supported different microbial loads and trajectories, implying that viral lysis modifies not only the physical properties of particles but also the composition and bioavailability of the released DOM. This interpretation aligns with studies showing that phytoplankton viral lysis generates DOM pools that are chemically distinct from healthy-diatom derived DOM and that these differences feed back onto bacterial succession, substrate use, and carbon cycling on and around aggregates^14,19,36^. Lastly, our results indicated that at least in this simplistic, culture-associated microbial community, natural senescence led to a similar community and functional shift as with viral infection, which has not been previously reported.^32^.

The strong response of Flavobacteriia (Bacteroidetes)—particularly *Polaribacter*— to physiological changes in both control and infected cultures, as revealed by our microscopy and omics data is consistent with the well-established role of *Polaribacter* and related Flavobacteria clades as bloom-associated specialists in the degradation of algal high-molecular-weight polysaccharides and late-bloom DOM ^37,38^;^39^. In our system, Bacteroidetes not only dominated the phycosphere but also proliferated on dead and decaying diatoms and subsequently became major components of the free-living community, matching observations that polysaccharide-degrading Bacteroidetes bridge attached and free-living niches during phytoplankton blooms ^40,41^. In contrast, both Alpha- and Gammaproteobacteria, which initially comprised ∼10% and ∼50% of the phycosphere, respectively, were consistently depleted from the phycosphere after viral infection. The stronger decline of Gammaproteobacteria in virus-treated cultures suggests that as substrates shifted from more labile to more structurally complex forms, copiotrophic lineages adapted to labile DOM were competitively displaced by polysaccharide specialists. The decrease in Alphaproteobacteria is consistent with their characterization as taxa that often thrive under more stable conditions, where they can act as vitamin suppliers, redox-balancing partners, and signalling hubs for phytoplankton—especially within the *Roseobacter* lineage ^8,42^. Viral disruption of host physiology would therefore be expected to destabilize niches typically occupied by Alpha- and many Gammaproteobacteria, while favouring Bacteroidetes that specialize in the breakdown of complex algal glycan^7,8,33,37,40^.

The DOM generated in this algal-virus system supported bacterial growth, in contrast to recent work on *Chaetoceros tenuissimus* ssRNA infected by an ssRNA virus, where virus-derived DOM did not sustain growth of *Sphingobacterium* isolates but stimulated their exopeptidase activity ^14^. Both systems involve Marnaviridae, but different genera (*Salisharnavirus* in our case versus *Bacillarnavirus* for *C. tenuissimus*). Together, our data suggests that the composition and bioavailability of viral-derived DOM depend on specific host–virus combinations. In our mixed microbial community, complementary enzymatic capabilities likely allowed coupling between hydrolysis and DOM assimilation: *Polaribacter* dominated carbohydrate active enzymes (CAZyme) expression, yet other Flavobacteriia also contributed (Figure 7,8), consistent with guild-level functional plasticity in flavobacterial CAZyme inventories during algal blooms ^38,41,43^. Despite under-representation of bacterial reads due to overwhelming viral RNA, metatranscriptomes still resolved clear functional shifts. Virus-treated samples showed marked upregulation of glycan-uptake machinery on Day 5—particularly TonB-dependent transporters and SusD-like proteins—relative to controls, in line with the use of these systems as proxies for polysaccharide scavenging during blooms ^44^. This indicates that transporter expression may be a more robust marker of bacterial glycan scavenging in high-viral-load contexts (Supplementary Figure 10). By Day 9, these differences diminished as control cultures entered senescence, suggesting that viral lysis primarily accelerated a successional trajectory that controls reached later under nutrient stress (Figure 9). Apparent incompleteness of some glycan-degradation pathways (e.g. βglucan, xylan, pectin) in individual samples most likely reflects coverage limitations rather than true pathway loss, whereas the enrichment of alginate degradation genes could be explained by the presence of alginate like or structurally analogous exopolysaccharides in diatoms ^7^.

### Host transcriptional reprogramming under viral dominance

Despite the overwhelming abundance of viral RNA, host-derived reads still provided insight into cellular reprogramming. The dominance of viral transcripts (∼99 % of mapped reads on Day 5) severely limited statistical power, and only a single host gene passed the adjusted-*p* threshold under the compositional ALDEx2 analysis. Traditional DESeq2 methods, which assume comparable total RNA counts across samples, yielded a larger set of differentially expressed genes but likely inflated fold changes due to compositional bias. ALDEx2’s CLR-based approach provided a conservative yet more reliable assessment under such skewed conditions. Even so, consistent expression patterns across infected replicates suggest that *Pseudo-nitzschia* cells mounted a stress response to infection. Upregulated transcripts included several heat-shock and stress-related genes including in the only WGCNA module significantly correlated with infection, indicative of oxidative stress responses like those described in other algal virus systems ^45^. In *Emiliania huxleyi* infections, for instance, antioxidant pathways such as glutathione metabolism are induced within 24–48 h ^45^. The concurrent increase in ribosomal protein transcripts in our dataset, although not statistically significant under ALDEx2, may indicate viral hijacking of the host translational machinery—a mechanism common among both DNA and RNA algal viruses ^46^. Conversely, transcripts associated with photosynthesis, silicon uptake, and central carbon metabolism were downregulated, reflecting the suppression of growth and biosynthetic activity typical of cells approaching lysis ^47^. Some of these shifts were also congruent between late control cultures and infected cultures, as evidenced by the *brown* module of the WGCNA analysis further suggesting that infection mimics senescence both in the host as well as the microbiome response. Together our observations portray a shift from active metabolism toward stress response and translational redirection under viral control.

### Experimental and analytical considerations

Our results also highlight the challenges of metatranscriptomic analysis under high viral loads. When viral transcripts dominate sequencing depth, even actively transcribed bacterial genes may appear depleted, and relative normalization methods can generate spurious differential expression. The clustering of predicted proteins before differential-expression testing improved statistical robustness but reduced resolution for specific carbohydrate-active enzymes—a trade-off that should be considered in future analyses. Early-stage sampling (Day 1 to 3 post infection) would capture infection onset before host RNA degradation and allow a clearer view of antiviral responses and the timing of bacterial activation. Moreover, RNA extraction from 1.2 µm filters captured primarily diatoms and particle-attached bacteria, while free-living taxa could be underrepresented, although the density and homogenization of the culture likely minimized this effect. Future experiments should include a complementary 0.22 µm fraction to assess the free-living community and disentangle its role in DOM turnover. Finally, the use of a long-term cultured diatom strain with an adapted microbiome may have reduced ecological variability compared to natural assemblages.

### Ecological implications and outlook

Together, our observations support a scenario in which PnGalRNAV infection precipitates premature senescence and lysis of *Pseudo-nitzschia*, releasing organic substrates that fuel heterotrophic bacterial growth. The accelerated recycling of carbon through the viral shunt demonstrates how viral activity can reshape microbial community function and shorten bloom turnover. However, not all diatom–virus systems behave identically; differences in viral host range, infection dynamics, and bacterial community composition may determine whether lysis primarily enhances DOM availability or promotes aggregation and export—the so-called viral shuttle ^16,18^. Preliminary unpublished results from our system, including limited aggregation potential in infected cultures ^48^, suggests that the PnGalRNAV–*Pseudo-nitzschia* interaction favours DOM production rather than aggregation.

Future studies comparing multiple diatom–virus pairs, sampled through the full infection cycle and coupled with metabolomic and physiological measurements, will be essential to determine whether the stress-related transcriptional changes and metabolic shutdown observed here represent universal features of algal virus infections or are system-specific outcomes.

## References

1 Seymour, J. R., Amin, S. A., Raina, J. B. & Stocker, R. Zooming in on the phycosphere: the ecological interface for phytoplankton-bacteria relationships. Nat Microbiol 2, 17065, doi:10.1038/nmicrobiol.2017.65 (2017).

2 Bell, W. & Mitchell, R. Chemotactic And Growth Responses Of Marine Bacteria To Algal Extracellular Products The Biological Bulletin 143, 10.2307/1540052 (1972).

3 Amin, S. A., Parker, M. S. & Armbrust, E. V. Interactions between diatoms and bacteria. Microbiol. Mol. Biol. Rev. 76, 667–684, doi:10.1128/MMBR.00007-12 (2012).

4 Bates, S. S., Douglas, D. J., Doucette, G. J. & Léger, C. Enhancement of domoic acid production by reintroducing bacteria to axenic cultures of the diatom *Pseudo-nitzschia multiseries*. Nat. Toxins 3, 428–435 (1995).

5 Kimura, K. & Tomaru, Y. Coculture with marine bacteria confers resistance to complete viral lysis of diatom cultures. Aquat. Microb. Ecol. 73, 69–80, doi:10.3354/ame01705 (2014).

6 Schafer, H., Abbas, B., Witte, H. & Muyzer, G. Genetic diversity of ‘satellite’ bacteria present in cultures of marine diatoms. FEMS Microbiol. Ecol. 42, 25–35, doi:10.1111/j.1574-6941.2002.tb00992.x (2002).

7 Ferrer-Gonzalez, F. X. et al. Resource partitioning of phytoplankton metabolites that support bacterial heterotrophy. ISME J 15, 762–773, doi:10.1038/s41396-020-00811-y (2021).

8 Buchan, A., LeCleir, G. R., Gulvik, C. A. & González, J. M. Master recyclers: features and functions of bacteria associated with phytoplankton blooms. Nature reviews. Microbiology 12, 686–698, doi:10.1038/nrmicro3326 (2014).

9 Teeling, H. et al. Substrate-controlled succession of marine bacterioplankton populations induced by a phytoplankton bloom. Science 336, 608–611, doi:10.1126/science.1218344 (2012).

10 Suttle, C. A. Marine viruses--major players in the global ecosystem. Nat. Rev. Microbiol. 5, 801–812, doi:10.1038/nrmicro1750 (2007).

11 Evans, C., Pearce, I. & Brussaard, C. P. Viral-mediated lysis of microbes and carbon release in the sub-Antarctic and Polar Frontal zones of the Australian Southern Ocean. Environ. Microbiol. 11, 2924–2934, doi:10.1111/j.1462-2920.2009.02050.x (2009).

12 Kuhlisch, C. et al. Viral infection of algal blooms leaves a unique metabolic footprint on the dissolved organic matter in the ocean. Sci Adv 7, doi:10.1126/sciadv.abf4680 (2021).

13 Diaz, B. P., Gallo, F., Moore, R. H. & Bidle, K. D. Virus infection of phytoplankton increases average molar mass and reduces hygroscopicity of aerosolized organic matter. Sci. Rep. 13, 7361, doi:10.1038/s41598-023-33818-4 (2023).

14 Kranzler, C. F. et al. Taxonomically distinct diatom viruses differentially impact microbial processing of organic matter. Sci Adv 11, eadq5439, doi:10.1126/sciadv.adq5439 (2025).

15 Zimmerman, A. E. et al. Metabolic and biogeochemical consequences of viral infection in aquatic ecosystems. Nat. Rev. Microbiol. 18, 21–34, doi:10.1038/s41579-019-0270-x (2020).

16 Laber, C. P. et al. Coccolithovirus facilitation of carbon export in the North Atlantic. Nat Microbiol 3, 537–547, doi:10.1038/s41564-018-0128-4 (2018).

17 Nissimov, J. I. et al. Dynamics of transparent exopolymer particle production and aggregation during viral infection of the coccolithophore, *Emiliania huxleyi*. Environ. Microbiol. 20, 2880–2897, doi:10.1111/1462-2920.14261 (2018).

18 Yamada, Y., Tomaru, Y., Fukuda, H. & Nagata, T. Aggregate Formation During the Viral Lysis of a Marine Diatom. Front. Mar. Sci. 5, doi:10.3389/fmars.2018.00167 (2018).

19 Xiao, X. et al. Viral Lysis Alters the Optical Properties and Biological Availability of Dissolved Organic Matter Derived from Prochlorococcus Picocyanobacteria. Appl. Environ. Microbiol. 87, doi:10.1128/AEM.02271-20 (2021).

20 Guillard, R. R. L. & Hargraves, P. E. Stichochrysis immobilis is a diatom, not a chrysophyte. Phycologia 32, 234–236, doi:10.2216/i0031-8884-32-3-234.1 (1993).

21 Turk Dermastia, T., Kutnjak, D., Gutierrez-Aguirre, I., Brussaard, C. P. D. & Bačnik, K. Discovery of novel and known viruses associated with toxigenic and non-toxigenic bloom forming diatoms from the Northern Adriatic Sea. Harmful Algae, 102737, 10.1016/j.hal.2024.102737 (2024).

22 Tinta, T. et al. Microbial Processing of Jellyfish Detritus in the Ocean. Front. Microbiol. 11, 590995, doi:10.3389/fmicb.2020.590995 (2020).

23 Glöckner, F. O. et al. An In Situ Hybridization Protocol for Detection and Identification of Planktonic Bacteria. Syst. Appl. Microbiol. 19, 403–406, 10.1016/S0723-2020(96)80069-5 (1996).

24 Bustin, S. A. et al. The MIQE guidelines: minimum information for publication of quantitative real-time PCR experiments. Clin. Chem. 55, 611–622, doi:10.1373/clinchem.2008.112797 (2009).

25 Parada, A. E., Needham, D. M. & Fuhrman, J. A. Every base matters: assessing small subunit rRNA primers for marine microbiomes with mock communities, time series and global field samples. Environ. Microbiol. 18, 1403–1414, doi:10.1111/1462-2920.13023 (2016).

26 McMurdie, P. J. & Holmes, S. phyloseq: An R Package for Reproducible Interactive Analysis and Graphics of Microbiome Census Data. PLoS One 8, e61217–e61217, doi:10.1371/journal.pone.0061217 (2013).

27. Oksanen, J., et al. (2019).

28 Charon, J., Murray, S. & Holmes, E. C. Revealing RNA virus diversity and evolution in unicellular algae transcriptomes. Virus Evolution 7, doi:10.1093/ve/veab070 (2021).

29 Wang, H. et al. Responses of Marine Diatom Skeletonema marinoi to Nutrient Deficiency: Programmed Cell Death. Appl. Environ. Microbiol. 86, doi:10.1128/AEM.02460-19 (2020).

30 Kirchman, D. L. & Kirchman, D. L. in Processes in Microbial Ecology 0 (Oxford University Press, 2018).

31 Sheik, A. R. et al. Responses of the coastal bacterial community to viral infection of the algae Phaeocystis globosa. ISME J 8, 212–225, doi:10.1038/ismej.2013.135 (2014).

32 Wilhelm, S. W. & Suttle, C. A. Viruses and Nutrient Cycles in the Sea aquatic food webs. Bioscience 49, 781–788 (1999).

33 Grossart, H.-P. & Ploug, H. Microbial degradation of organic carbon and nitrogen on diatom aggregates. Limnol. Oceanogr. 46, 267–277, doi:10.4319/lo.2001.46.2.0267 (2001).

34. (!!! INVALID CITATION !!! 1,34,35).

35 Kiorboe, T., Tang, K., Grossart, H. P. & Ploug, H. Dynamics of microbial communities on marine snow aggregates: colonization, growth, detachment, and grazing mortality of attached bacteria. Appl. Environ. Microbiol. 69, 3036–3047, doi:10.1128/AEM.69.6.3036-3047.2003 (2003).

36 Lønborg, C., Middelboe, M. & Brussaard, C. P. D. Viral lysis of *Micromonas pusilla*: impacts on dissolved organic matter production and composition. Biogeochemistry 116, 231–240, doi:10.1007/s10533-013-9853-1 (2013).

37 Xing, P. et al. Niches of two polysaccharide-degrading Polaribacter isolates from the North Sea during a spring diatom bloom. ISME J 9, 1410–1422, doi:10.1038/ismej.2014.225 (2015).

38 Avci, B., Kruger, K., Fuchs, B. M., Teeling, H. & Amann, R. I. Polysaccharide niche partitioning of distinct Polaribacter clades during North Sea spring algal blooms. ISME J 14, 1369–1383, doi:10.1038/s41396-020-0601-y (2020).

39 Redondo-Rio, A., Mundy, C. J., Tamames, J. & Pedros-Alio, C. Specialized Bacteroidetes dominate the Arctic Ocean during marine spring blooms. Front. Microbiol. 15, 1481702, doi:10.3389/fmicb.2024.1481702 (2024).

40 Wang, F. Q. et al. Particle-attached bacteria act as gatekeepers in the decomposition of complex phytoplankton polysaccharides. Microbiome 12, 32, doi:10.1186/s40168-024-01757-5 (2024).

41 Ma, K.-J. et al. Polysaccharide metabolic pattern of Cytophagales and Flavobacteriales: a comprehensive genomics approach. Front. Mar. Sci. 12, doi:10.3389/fmars.2025.1551618 (2025).

42 Geng, H. & Belas, R. Molecular mechanisms underlying roseobacter-phytoplankton symbioses. Curr. Opin. Biotechnol. 21, 332–338, doi:10.1016/j.copbio.2010.03.013 (2010).

43 Li, X. et al. Dynamic patterns of carbohydrate metabolism genes in bacterioplankton during marine algal blooms. Microbiol. Res. 286, 127785, doi:10.1016/j.micres.2024.127785 (2024).

44 Francis, T. B. et al. Changing expression patterns of TonB-dependent transporters suggest shifts in polysaccharide consumption over the course of a spring phytoplankton bloom. ISME J 15, 2336–2350, doi:10.1038/s41396-021-00928-8 (2021).

45 Rosenwasser, S. et al. Rewiring Host Lipid Metabolism by Large Viruses Determines the Fate of Emiliania huxleyi, a Bloom-Forming Alga in the Ocean. Plant Cell 26, 2689–2707, doi:10.1105/tpc.114.125641 (2014).

46 Thomy, J., Schvarcz, C. R., McBeain, K. A., Edwards, K. F. & Steward, G. F. Eukaryotic viruses encode the ribosomal protein eL40. Npj Viruses 2, 51, doi:10.1038/s44298-024-00060-2 (2024).

47 Kranzler, C. F. et al. Silicon limitation facilitates virus infection and mortality of marine diatoms. Nat Microbiol 4, 1790–1797, doi:10.1038/s41564-019-0502-x (2019).

48. Turk Dermastia, T., et al. in 12th Aquatic Virus Workshop. (ed Sheree Yau).

