## Supplementary Methods for "Viral lysis of a toxigenic diatom triggers a microbial response mimicking hastened senescence"

Raw metatranscriptomic reads were quality-checked and adapter-trimmed using *CLC Genomics Workbench* (QIAGEN) with default parameters for RNA-Seq workflows. High-quality paired-end reads were then assembled de novo using *rnaSPAdes* (v3.15.4), optimized for transcriptome reconstruction without a reference genome ^1^. The assembled transcriptome was translated using *TransDeoder* (v5.7.1) (<https://github.com/TransDecoder/TransDecoder>) and the resulting peptides were clustered at 98% using CD-HIT. The top transcript in each cluster was used as a reference index for read mapping. Transcript-level abundance was quantified per sample using *salmon* (v1.9.0) in quasi-mapping mode ^2^. For functional and taxonomic annotation, the full transcriptome was run through *eggNOG-mapper* (v2.1.12) using *DIAMOND*-based searches against the eggNOG database with the --itype CDS mode ^3^. Annotated transcripts were assigned NCBI TaxIDs, which were then mapped to higher-order taxonomic domains (e.g., Bacteria, Archaea, Eukaryota, Viruses) by querying the eggNOG SQLite taxonomy database and recursively tracing parent lineages to established domain-level taxIDs. This domain-level classification was merged with sample-level transcript abundances to produce per-sample summaries of transcriptional activity across major biological domains, enabling downstream ecological and functional comparisons of microbial and host community dynamics.

To assess the taxonomic composition of the bacterial community present in the metatranscriptome, we used *Kraken2* **(v2.1.5)** with the *--standard* configuration ^4^, comprising complete genomes from archaea, bacteria, and viruses available in RefSeq, as well as associated taxonomy files. Interleaved paired-end reads were processed by Kraken2 in single-end mode, as Kraken2 does not support interleaved input. Taxonomic summaries were generated at the genus and family levels to compare with the 16S amplicon analysis. Only transcripts annotated as bacterial were retained for downstream differential expression (DE) analysis.

To account for compositional constraints inherent to metatranscriptomic data, *ALDEx2* (v1.30.0) was used to identify differentially expressed bacterial genes between treatments ^5^. Salmon-derived transcript counts were extracted and filtered to include only bacterial transcripts, and centered log-ratio (CLR) transformation was performed using aldex.clr. Differential abundance testing was conducted using Welch’s t-test (aldex.ttest) with Benjamini–Hochberg FDR correction. Genes with effect sizes >1 and adjusted *p*-values (*eBH*) < 0.1 were considered significantly differentially expressed.

To gain functional insight into transcripts lacking annotation in the eggNOG-mapper pipeline, we submitted unannotated but significantly differentially expressed sequences to *AlphaFold2* via *ColabFold* ^6^ and performed structure-based homology searches using *FoldSeek* ^7^ against the AlphaFoldDB. Hits were manually curated and used to update functional annotations of previously uncharacterized genes. These were integrated into downstream analyses, including pathway enrichment and genus-function association visualizations.

To estimate the completeness of microbial carbohydrate utilization pathways, peptides obtained with *TransDecoder* on the entire set of assembled transcripts, were further annotated with dbCAN3 ^8^. We curated a set of marker CAZy families ^9^ and PUL-associated genes representative of seven polysaccharide substrates or substrate analogues that are produced by diatoms: β-glucan (laminarin), alginate, chitin, xylan, β -mannan, exopolysaccharide and host-glycan. Genes and gene families associated with degradation of these substrates were sourced from the dbCANsub database. Pathway completeness per sample was calculated as the proportion of these marker families detected (at least one transcript with TPM > 0.1), divided by the total number of families found in our data across all samples for each substrate. We are aware this is less conservative estimate than using the entire set of known CAZy families but it is unlikely that a selected set of bacteria present in the cultures would possess all the different degradation enzymes. Importantly, we also extended traditional CAZy classification by including non-enzymatic but functionally essential transport and binding proteins, namely SusD-like and TonB-dependent receptors, based on PFAM and eggNOG annotations. This approach reflects the modular organization of Polysaccharide Utilization Loci (PULs) in marine bacteria typically used in the analysis of metagenomes and helps capture functional shifts in glycan targeting strategies even when classical enzymatic markers are underrepresented. We acknowledge that this is a minimalist marker set, and additional CAZy families (e.g., auxiliary debranching enzymes, additional CBMs, or alternative lyases) likely contribute to full pathway function. However, this conservative definition offers a standardized and interpretable framework for comparing pathway erosion or expansion across treatments.

Host transcriptome

We analysed the diatom host transcriptomic response to viral infection using a reference-based approach leveraging publicly available *Pseudo-nitzschia* transcriptomes from the MMETSP project ^10^. These transcriptomes were concatenated and dereplicated using *CD-HIT* ^11^ and functionally annotated with dammit, which integrates tools including *TransDecoder*, *hmmer* (v3.4) ^12^*,* and *diamond* (v2.1.11) ^13^. Transcript quantification was performed using s*almon*, aligning metatranscriptomic reads from each sample to the annotated reference transcriptome. TPM values were extracted and used for expression analysis across all samples. To assess DE, we first used *ALDEx2* as for the bacterial metatranscriptomes, but this analysis was unsatisfactory so we used *DESeq2* (v1.38.2) ^14^ as well. The experimental conditions were *virus5* vs. *control5* and diatom biomass as a covariate. Annotation of differentially expressed transcripts was enhanced using *eggNOG-mapper*, providing COG, KEGG, and CAZy identifiers. To evaluate host-transcriptome completeness, we performed a completeness analysis using the Benchmarking Universal Single-Copy Orthologs (BUSCO) database (v. 5.7.1) ^15^ using the stremenopile database subset as reference.

We focused on interpreting the response of the diatom host, emphasizing transcripts associated with photosynthesis, translation, silicon metabolism, and TEP-related carbohydrate production. TPM expression matrices were merged with DE results to generate heatmaps and identify functional trends. CAZy-annotated genes and those involved in KEGG pathways for sugar metabolism (e.g., ko00520, ko00500) were further examined. We assessed pathway completeness and visualized host metabolic pathways using KEGG Mapper and Pathview, integrating both gene presence and DE patterns. Overall, this analysis allowed us to track broad-scale host metabolic shifts and identify stress-responsive and potentially virus-targeted functional modules.

Variance-stabilized expression values were obtained from the full host transcriptome by applying the DESeq2 vst() transformation to filtered count data (genes with nonzero counts in at least one sample). Transcripts previously annotated with PFAM, KEGG Orthologs (KO), and COG categories were used to define gene sets of interest (e.g., stress-related or ATPase-associated). For visual exploration, TPM-normalized values (from *tximport*) were plotted across all samples as Z-score-scaled heatmaps using the *pheatmap* R package. Transcripts significantly differentially expressed (adjusted p-value < 0.05, |log₂FC| > 1) in the virus5 vs control5 comparison were labeled by direction of change.

To evaluate broader transcriptional patterns and co-expression a network analysis was performed using the WGCNA R package. The input matrix consisted of variance-stabilized expression values of transcripts expressed in at least one sample. A soft-thresholding power was chosen based on scale-free topology fitting. Genes were clustered using hierarchical clustering of the topological overlap matrix, and modules were identified via dynamic tree cutting. Module eigengenes were correlated with sample traits including viral condition and diatom biomass. Modules containing heat shock or HATPase-associated transcripts were extracted for further inspection. KEGG-based functional enrichment of selected modules (e.g., the yellow module containing several HATPase genes) was performed using the clusterProfiler package. KEGG Orthologs associated with module transcripts were extracted and used to build custom gene sets for enrichment testing. Dotplots were generated to visualize statistically enriched pathways, with adjusted p-values (FDR < 0.1) used to define significance.

1	Bushmanova, E., Antipov, D., Lapidus, A. & Prjibelski, A. D. rnaSPAdes: a de novo transcriptome assembler and its application to RNA-Seq data. *Gigascience* **8**, doi:10.1093/gigascience/giz100 (2019).

2	Patro, R., Duggal, G., Love, M. I., Irizarry, R. A. & Kingsford, C. Salmon provides fast and bias-aware quantification of transcript expression. *Nat. Methods* **14**, 417-419, doi:10.1038/nmeth.4197 (2017).

3	Cantalapiedra, C. P., Hernandez-Plaza, A., Letunic, I., Bork, P. & Huerta-Cepas, J. eggNOG-mapper v2: Functional Annotation, Orthology Assignments, and Domain Prediction at the Metagenomic Scale. *Mol. Biol. Evol.* **38**, 5825-5829, doi:10.1093/molbev/msab293 (2021).

4	Wood, D. E., Lu, J. & Langmead, B. Improved metagenomic analysis with Kraken 2. *Genome Biol.* **20**, 257, doi:10.1186/s13059-019-1891-0 (2019).

5	Fernandes, A. D., Macklaim, J. M., Linn, T. G., Reid, G. & Gloor, G. B. ANOVA-like differential expression (ALDEx) analysis for mixed population RNA-Seq. *PLoS One* **8**, e67019, doi:10.1371/journal.pone.0067019 (2013).

6	Mirdita, M. *et al.* ColabFold: making protein folding accessible to all. *Nat. Methods* **19**, 679-682, doi:10.1038/s41592-022-01488-1 (2022).

7	van Kempen, M. *et al.* Fast and accurate protein structure search with Foldseek. *bioRxiv*, doi:10.1101/2022.02.07.479398 (2023).

8	Zheng, J. *et al.* dbCAN3: automated carbohydrate-active enzyme and substrate annotation. *Nucleic Acids Res.* **51**, W115-W121, doi:10.1093/nar/gkad328 (2023).

9	Cantarel, B. L. *et al.* The Carbohydrate-Active EnZymes database (CAZy): an expert resource for Glycogenomics. *Nucleic Acids Res.* **37**, D233-238, doi:10.1093/nar/gkn663 (2009).

10	Keeling, P. J. *et al.* The Marine Microbial Eukaryote Transcriptome Sequencing Project (MMETSP): illuminating the functional diversity of eukaryotic life in the oceans through transcriptome sequencing. *PLoS Biol.* **12**, e1001889, doi:10.1371/journal.pbio.1001889 (2014).

11	Li, W. & Godzik, A. Cd-hit: a fast program for clustering and comparing large sets of protein or nucleotide sequences. *Bioinformatics* **22**, 1658-1659, doi:10.1093/bioinformatics/btl158 (2006).

12	Eddy, S. R. Accelerated Profile HMM Searches. *PLoS Comput. Biol.* **7**, e1002195, doi:10.1371/journal.pcbi.1002195 (2011).

13	Buchfink, B., Reuter, K. & Drost, H. G. Sensitive protein alignments at tree-of-life scale using DIAMOND. *Nat. Methods* **18**, 366-368, doi:10.1038/s41592-021-01101-x (2021).

14	Love, M. I., Huber, W. & Anders, S. Moderated estimation of fold change and dispersion for RNA-seq data with DESeq2. *Genome Biol.* **15**, 550, doi:10.1186/s13059-014-0550-8 (2014).

15	Manni, M., Berkeley, M. R., Seppey, M., Simao, F. A. & Zdobnov, E. M. BUSCO Update: Novel and Streamlined Workflows along with Broader and Deeper Phylogenetic Coverage for Scoring of Eukaryotic, Prokaryotic, and Viral Genomes. *Mol. Biol. Evol.* **38**, 4647-4654, doi:10.1093/molbev/msab199 (2021).
