## Supplementary Table 11 for "Viral lysis of a toxigenic diatom triggers a microbial response mimicking hastened senescence"

**Supplementary Table 11. Functional summary of WGCNA modules and top enriched KEGG Orthologs (KOs).** Arrows indicate the direction of eigengene–trait correlations (↑ positive, ↓ negative, ↔ weak/no correlation). Asterisks (*) denote significant correlations ($p < 0.05$). KO terms are based on enrichment of module transcripts against KEGG Orthology annotations.

| Module | Correlation pattern | Functional summary | Representative enriched KOs (from enrichment files) |
|---|---|---|---|
| **Black** | **↑* Virus5, ↑* Virus9, ↓ Controls and Initial** | **Glyoxylate cycle and stress response.** *Note: Most genes annotated as Gammaproteobacteria (likely bacterial origin).* | K01637 Isocitrate lyase; K01638 Malate synthase; K03088 ECF sigma factor; K02406 Flagellin C-term; K03969 Thylakoid organization; K21841 Anti-nitrosylation; K03283 Heat-shock protein; K03704 Cold-shock domain. |
| **Brown** | **↑ Virus5, ↓* Virus9, ↑ Control5, ↓ Control9, ↑* Initial** | **Chromatin, translation and energy metabolism.** | K11275 Histones H2A/H2B/H3/H4; K02983 Ribosomal eS30; K01100 Fructose-1,6-bisphosphatase N-term; K00248 Acyl-CoA dehydrogenase; K03521 ETF domain; K12580 NOT complex subunit. |
| **Turquoise** | **↓ Virus5, ↓ Virus9, ↑ Control5, ↔ Control9, ↑* Initial,** | **Translation and photosynthetic core.** | K00134 GAPDH; K04077 HSP60 (chaperonin); K02358 EF-Tu; K02641 Oxidoreductase NAD-binding; K01624 FBA class II; K01915 Glutamine synthetase; K00789 SAM synthase; K02716 PSII Mn-stabilizing protein; K04567 tRNA synthetase II. |
| **Magenta** | **↑ Virus5, ↓ Virus9, ↔ Controls and Initial** | **Membrane and ER remodeling.** | K00814 Alanine aminotransferase; K13989 ERAD component; K05643/45 ABC transporters; K11262 Acetyl-CoA carboxylase; K00731 Polypeptide GalNAc-transferase; K02183 EF-hand $Ca^{2+}$ motif; K13412 LPS kinase-like family. |
